# Tau conveys intrinsic hyperactivity of VTA dopamine neurons but an inability to sustain burst firing

**DOI:** 10.1101/2025.07.28.666953

**Authors:** William M. Kennedy, Eva Troyano-Rodriguez, Matthew H. Higgs, Harris E. Blankenship, Anu Korukonda, David Weinshenker, Heather C. Rice, Michael J. Beckstead

## Abstract

**INTRODUCTION:** Ventral tegmental area (VTA) dopamine has been implicated in neuropsychiatric symptoms observed in Alzheimer’s disease (AD) patients. Dopaminergic dysfunction and aberrant firing are observed in mouse AD models, but the specific roles of Aβ and tau have not been determined.

**METHODS:** We performed electrophysiological recordings of single VTA dopamine neuron firing in the 3xTg-AD model, followed by recordings in amyloid (APP^NL-G-F^)- and human tau (hTau)-based models to determine the pathological triggers of impaired firing.

**RESULTS:** *In vivo* dopamine neuron recordings showed fewer spikes in defined bursts in 3xTg-AD mice versus controls. *Ex vivo* studies showed an impaired ability to sustain firing during depolarization, which was mimicked with depolarized current in wild type neurons. Dopamine neurons transduced with hTau reflected firing aberrations and impaired bursting, but the effects were not recapitulated in the APP^NL-G-F^ model,

**DISCUSSION:** These results suggest that hTau specifically induces hyperexcitable states within individual dopamine neurons, disrupting burst firing. This dopaminergic dysfunction could compromise reward learning and contribute to the psychiatric symptoms observed in AD.

## 1 INTRODUCTION

Despite recent advances, highly efficacious treatments for Alzheimer’s disease (AD) remain elusive, likely due to a lack of understanding about early stages of pathology that occur prior to neurodegeneration. While the pronounced learning and memory deficits associated with AD correlate with hippocampal degeneration, psychiatric symptoms including anxiety and depression precede, or appear simultaneously with, memory and cognitive impairment [1-3]. Ventral tegmental area (VTA) dopamine neurons play prominent roles in anxiety and depression phenotypes, suggesting involvement in AD-related psychopathology. Work by us and others has described altered dopaminergic firing, decreased dopamine release, and loss of dopaminergic neurons in mouse models of AD [4, 5], providing a plausible link between dopamine disruption and the onset of AD-like phenotypes. Recently, we described dopaminergic hyperactivity in brain slices from the 3xTg-AD mouse model that combines amyloid beta (Aβ) and hyperphosphorylated human tau (hTau) pathology [5]. Specifically, 3xTg-AD dopamine neurons exhibited faster firing and disrupted pacemaking that was due to casein kinase 2 (CK2)-dependent disruption of small conductance calcium-activated potassium (SK) channel currents [5]. However, it is not known how dopaminergic firing is affected *in vivo* in AD mouse models and whether tau or amyloid, or both, is responsible for the hyperactivity.

VTA dopamine neurons project to a broad array of brain regions that are important for anxiety, learning and memory, aversion, and reward [3, 6-10]. We previously found that 3xTg-AD mice show impaired dopamine-dependent reward learning by 6 months of age [5]. Reward learning requires phasic, time sensitive release of dopamine in key projection areas [11-15]. Thus, impaired burst firing may be a contributing factor in disrupted reward learning in 3xTg-AD mice. Normally, excitatory input readily induces burst firing in dopamine neurons; however, dopamine neurons are also susceptible to voltage-gated sodium channel block during prolonged depolarization [16-18] that may limit the duration of bursts. Consistent with this possibility, VTA dopamine neurons from 3xTg-AD mice exhibit slightly depolarized action potential thresholds, suggesting potential inactivation of voltage-gated sodium channels [5].

Here, we sought a better understanding of the hyperexcitability observed in VTA dopamine neurons from AD mice by investigating the specific roles of Aβ and tau pathology. In addition to 3xTg-AD mice, we used APP^NL-G-F^ knock-in mice to model amyloid accumulation under the amyloid precursor protein’s endogenous promoter [19, 20] and also used a virus-induced overexpression of hyperphosphorylated hTau specifically in VTA dopamine neurons. We found that 3xTg-AD dopamine neurons *in vivo* showed fewer spikes in defined bursts, which might result from a depolarization-induced inability to sustain firing. Further, we found that dopamine neurons expressing hTau, but not dopamine neurons from the APP^NL-G-F^ model, exhibited fast, irregular firing and cessation of prolonged bursts. Our results suggest that hTau rather than Aβ is responsible for the disruption of intrinsic dopamine neuron firing in AD models.

## 2 METHODS

### 2.1 Animals

Three models of AD were used. 3xTg-AD mice (MMRRC #034830) and non-transgenic controls on a hybrid background (C57BL/6; 129×1/SvJ; 129S1/Sv embryo injected in a B6;129) were obtained from Dr. Salvadore Oddo [20]. 3xTg-AD mice carry three different transgenes with mutations relevant to AD: PSEN1_M146V_, APP_Swe_, and Tau_P301L_. We used 13-15-month-old female mice for *in vivo* studies and 13-15-month-old male and female mice for *ex vivo* recordings. We also used 6-month-old male and female APP^NL-G-F^ mice, which carry three mutations to the APP gene: APP_Swe_, APP_1716F_, and APP_E693G_ [19]. Because of their background, C57BL/6 mice were used as controls for APP^NL-G-F^ experiments.The third AD model was achieved with bilateral stereotaxic injections of an AAV construct encoding a hyperphosphorylated, mutated human tau (AAV5-DIO-hTau_P364S_; 300 nL, 2.5 × 10^13^ GC/mL) directly into the VTA (AP −2.91 mm, ML ±0.70 mm, and DV −4.75 mm) of 5-month-old DAT^Cre^ mice (JAX#006660). DAT^Cre^ mice with or without AAV5-DIO-eYFP bilateral stereotaxic injections were used as controls. All Injected mice were ∼5-months-old and were used at ∼6-months, 3-6 weeks post-injection. All *ex vivo* studies showed no indication of sex differences, and therefore data were pooled. All mice were group-housed in ventilated standard cages with ad libitum access to food and water. Animal rooms were maintained on a 12-h light/dark cycle (3xTg, 3xTg-CTR, C57BL/6 lights off at 9:00 AM; DAT^Cre^, APP^NL-G-F^ lights on at 7:00 AM) with the temperature held at ∼26°C. Procedures were in accordance with the Guide for the Care and Use of Laboratory Animals and were approved by the Institutional Animal Care and Use Committee at the Oklahoma Medical Research Foundation (OMRF). Studies were conducted in compliance with ARRIVE guidelines.

### 2.2 *In vivo* electrophysiology

For *in vivo* recordings, mice were deeply anesthetized with chloral hydrate (initial dose 400 mg/kg, i.p.) and were positioned in a stereotaxic apparatus (Kopf Instruments, Tujunga, California). The plane of anesthesia was continually monitored by absence of eye blink to a saline drop or any response to a gentle toe pinch. When necessary, a supplemental dose of 50-70 mg/kg was given i.p. with a syringe pump. Body temperature was maintained at 35 - 36.5°C using a heating pad (SomnoSuite, Kent Scientific corporation). Once the skull was exposed, a hole was drilled at the appropriate coordinates as described previously [21]. Polished glass micropipettes with an internal diameter of 1.5 mm (Sutter Instruments, Novato, California) were pulled with a PC-10 puller (Narishige International) to a shank length of ∼1 cm. Pipettes were then filled with 2% pontamine sky blue dye in 2 M sodium acetate (pH 7.5), and were cut to a final resistance of 5-9 MΩ (Glass electrode R/C meter, model 2700, A-M systems; Carlsborg, WA). The electrode was advanced into the brain using a hydraulic microdrive (Stoelting, Wood Dale, Illinois). The VTA was localized using stereotaxic coordinates according to the Paxinos and Franklin mouse brain atlas [22]: AP −2.9 to −3.6 mm, ML 0.4 to 0.6 mm, and DV: −3.8 to −5 mm. One to six electrode tracks were run in each mouse. Single-unit recordings were obtained using a MultiClamp 700B amplifier (Molecular Devices). The initial signal was amplified and then filtered within a bandpass of 0.1-2.0 kHz. The data were collected with Axograph X and later analyzed offline using custom Python scripts.

Putative dopamine neurons were identified based on established electrophysiological properties: a broad action potential (> 1.8 ms), bi- or triphasic waveform, and slow firing (1-10 Hz) [21, 23, 24], followed by histological verification of the recording track location in the VTA. A gaussian filter was used to minimize noise prior to spike detection. For each cell, spike activity was collected over at least 100 spikes spanning 100-second to 10-minute recordings. The coefficient of variation of the inter-spike interval (CV_ISI_) was calculated as the standard deviation of the ISI divided by the mean.Burst activity of dopamine neurons was analyzed based on published criteria [18]. A burst was defined as a train of at least two spikes with an initial ISI of ≤ 80 milliseconds and an ISI of >160 milliseconds defined the end of the burst [18]. The parameters analyzed included the percentage of spikes fired in bursts, the mean burst length, the mean burst ISI, and the length of the first intra-burst ISI. Non-burst ISIs were separated by exclusion from the bursts and analyzed for spike frequency and CV_ISI_. On the final track for each mouse, pontamine sky blue dye was injected iontophoretically as described previously [21, 24]. Mice were then sacrificed, and their brains removed and placed in 4% paraformaldehyde for 1-4 days. Localization of the recording site was made by visual inspection of a pontamine dot in 80 μm coronal brain sections.

### 2.3 *Ex vivo* electrophysiology

Mice were anesthetized with isoflurane and sacrificed by decapitation. After quick brain removal, horizontal sections (200 μm) containing the ventral midbrain were sectioned on a vibrating microtome (Leica VT1200S). The cutting solution was a choline chloride-based solution containing the following (in mM): 110 choline chloride, 2.5 KCl, 1.25 NaH_2_PO_4_, 0.5 CaCl_2_, 7 MgSO_4_, 25 glucose, 26 NaHCO_3_,11.6 Na ascorbate, and 3.1 Na pyruvate. Ice-cold cutting solution was bubbled with 95% O_2_, 5% CO_2_. Slices were then allowed to recover at 35-37°C for 30 minutes in artificial cerebrospinal fluid (aCSF) containing the following (in mM): 126 NaCl, 2.5 KCl, 1.2 MgCl_2_, 2.4 CaCl_2_, 1.2 NaH_2_PO_4_, 21.4 NaHCO3, and 11.1 glucose, bubbled with 95% O_2_, 5% CO_2_. Slices were then moved to room temperature for a minimum of 30 minutes, and taken for recording up to 8 hours after preparation. Slices were transferred to a recording chamber where they were perfused with warmed aCSF (Warner Instruments inline heater, 31 ± 2°C) at a rate of 2-3 mL/min via gravity or a peristaltic pump (Warner Instruments). Slices were visualized under Dödt gradient contrast optics via an upright microscope (Nikon).

Patch clamp pipettes were pulled from thick-walled borosilicate glass with filament (1.5/0.86 mm; Warner Instruments). Perforated-patch whole-cell recordings used the gramicidin perforating agent with 2-5 MΩ pipettes filled with the following (in mM): 145.5 KCl, 7.5 NaCl, 10 HEPES, pH 7.0, and 2 μg/ml gramicidin-D (MP Biomedicals). A stock solution of gramicidin-D in dimethylsulfoxide was diluted in the pipette solution and vortexed for ∼30 s to achieve a final working concentration (2-5 µg/mL). If the gramicidin-containing pipette solution clogged the pipette, the stock solution was sonicated for 5-10 minutes before dilution. The pipette tip was filled with the same solution without gramicidin to prevent interference of seal formation. After forming a seal on the cell membrane, perforation was allowed to proceed until adequate bridge balance and capacitance neutralization were obtained before starting experiments. Series resistance (R_series_) was monitored throughout experiments with small current steps and corrected using bridge balance. R_series_was under 100 MΩ for all experiments and not different between compared groups. Experiments were discarded if R_series_ changed rapidly, indicating breakage of the perforated patch.

Recordings were acquired with a MultiClamp 700B amplifier and digitized with an Instrutech ITC-18 board. AxoGraph X and LabChart (ADInstruments) software were used for recording. Current-clamp recordings were low-pass filtered at 10 kHz and digitized at 20 kHz. VTA dopamine neurons were identified first by location (>100 μm medial to the medial terminal nucleus of the accessory optic tract near the level of the fasciculus retroflexus). Electrophysiological criteria for VTA dopamine neuron identification included slow (0.5-8 Hz) spontaneous firing in the cell-attached configuration prior to perforation and wide (>1 ms) spikes [25], the presence of a rebound delay after a hyperpolarizing step (−100 pA) [26, 27], and/or presence of an A-type potassium current in voltage clamp. These criteria were previously used to identify dopamine neurons in gramicidin-perforated patch recordings [5, 27]. For the hTau experiment, eYFP epifluorescence was used to identify control dopamine neurons and no differences of intrinsic properties were observed with or without virus injections. ISI trajectories were analyzed as previously described, with the addition of 95% confidence intervals estimated in both the x (time) and y (voltage) coordinates [5]. For analysis of spontaneous firing, action potentials were binned into 10 s windows and CV_ISI_was measured in each bin and averaged across bins to minimize the contribution of firing rate drift. Frequency-current (F-I) curves were constructed after a stable baseline of intrinsic firing was established. Current injection steps of 25 pA from -100 to +200 pA were 2 s long with 10 s from start to start. F-I curves were analyzed in Axograph X with an event detection threshold of 3-5 mV/ms and visually checked to ensure all events were captured and erroneous events (noise spikes, synaptic events, etc.) were discarded. Current steps were segmented into two bins (0.01-1.00 s and 1.01-2.00 s to minimize erroneous capture of current steps and double capture of spikes) for analysis of spike rate adaptation.

### 2.4 Immunofluorescence

Methoxy-X04 was diluted in DMSO and sterile saline to a working solution of 2mg/ml, which was injected 3 hours prior to brain collection. Mice were first anesthetized by inhalation of isoflurane and deeply anesthetized with subsequent intraperitoneal injection of 2,2,2-tribromoethanol (Avertin;Sigma-Aldrich). Anesthetized mice were perfused intracardially with 0.1 M PBS and chilled 4% PFA before brains were removed and postfixed overnight in PFA. Free-floating horizontal slices were made on a Leica VT1200S (50 μm). For posthoc staining of phosphorylated tau and tyrosine hydroxylase, slices were permeabilized with 0.4% PBST (Triton X-100) and blocked with 7% normal donkey serum in PBST. Primary antibodies were diluted as follows: chicken anti-tyrosine hydroxylase (TH, 1:1000, Abcam), rabbit anti-phosphorylated tau (pTau, 1:500) and incubated together for 48 hours at 4°C. Secondary antibodies were diluted as follows: anti-chicken AlexaFluor 488 (1:200, JacksonImmuno), anti-rabbit AlexaFluor 594 Plus (1:500, Invitrogen) and incubated together at room temperature for 2 hours. Slices were mounted with ProLong Gold (Invitrogen). Slices were imaged on a Zeiss LSM-710 confocal microscope with a 10× objective for tiling or 40x for single neuron visualization using a z-step of 5 μm.

### 2.5 Chemical reagents

Chloral hydrate (Sigma-Aldrich) was dissolved in sterile saline (0.9%, Hospira Inc.). Pontamine sky blue dye was obtained from Alfa Aesar. All other salts and chemicals were purchased from Fischer Scientific or Sigma-Aldrich.

### 2.6 Statistical analysis

Data are expressed as mean ± SEM. All tests used a two-tailed Type I error rate (α) of 0.05 (GraphPad Prism, La Jolla, CA). We used a 1-way repeated measures (RM) ANOVA with Tukey’s multiple comparison in Figure 3. 2-way RM ANOVAs with Bonferroni’s multiple comparisons were used to assess genotype or S1-S2 differences. Interaction effects were reported, but no post-hoc analysis were done to avoid false discoveries from numerous comparisons. Data were tested for the assumptions of the respective comparisons. If assumptions were not met, the data were further investigated for outliers. If data were non-normally distributed or had significantly different variances, nonparametric tests were used (e.g., Mann-Whitney U for nonparametric tests with 2 groups). All t-tests were 2 sided. DF, F, W, U, p, and n values are indicated in the text, figures, or figure legends.

## 3 RESULTS

### 3.1 *In vivo* firing of dopamine neurons in the 3xTg-AD mouse model

We first made *in vivo* recordings of putative dopamine neurons in the VTA of wild type and 3xTg-AD mice. We recorded from the VTA of deeply anesthetized mice in a stereotaxic apparatus (**Fig.1A**). Dopamine neurons were identified based on their action potential waveform (spike width > 1.8 ms, **Fig. 1B**). We identified individual spikes by threshold detection (see 2.2) and calculated the overall firing rate and the ISI for each recorded neuron. The localization of recording tracks within the VTA was confirmed by pontamine labelling at the end of each experiment (**Fig. 1C**). Firing rates were consistent with previous *in vivo* reports dopamine neurons (<10 Hz in anesthetized rodents) [23].Recorded neurons revealed no overall difference in firing rate between wild type and 3xTg-AD mice (3xTg 4.0 ± 0.3 Hz; WT: 4.2 ± 0.4 Hz; **Fig. 1D**). However, the coefficient of variation of the ISI (CV_ISI_), a measure of the rhythmicity of spiking, was lower (i.e., more rhythmic) in VTA dopamine neurons from 3xTg-AD mice than wild type (3xTg: 0.51 ± 0.07; WT: 0.74 ± 0.04; **Fig. 1D**). This was unexpected, given our previous results obtained in brain slices that suggested overall hyperactivity and more irregularity in VTA dopamine neurons of the 3xTg-AD model [5]. However, dopamine neurons in previous studies have shown large differences in firing rates and patterns *in vivo* compared to *ex vivo* [28].

**Figure 1.**
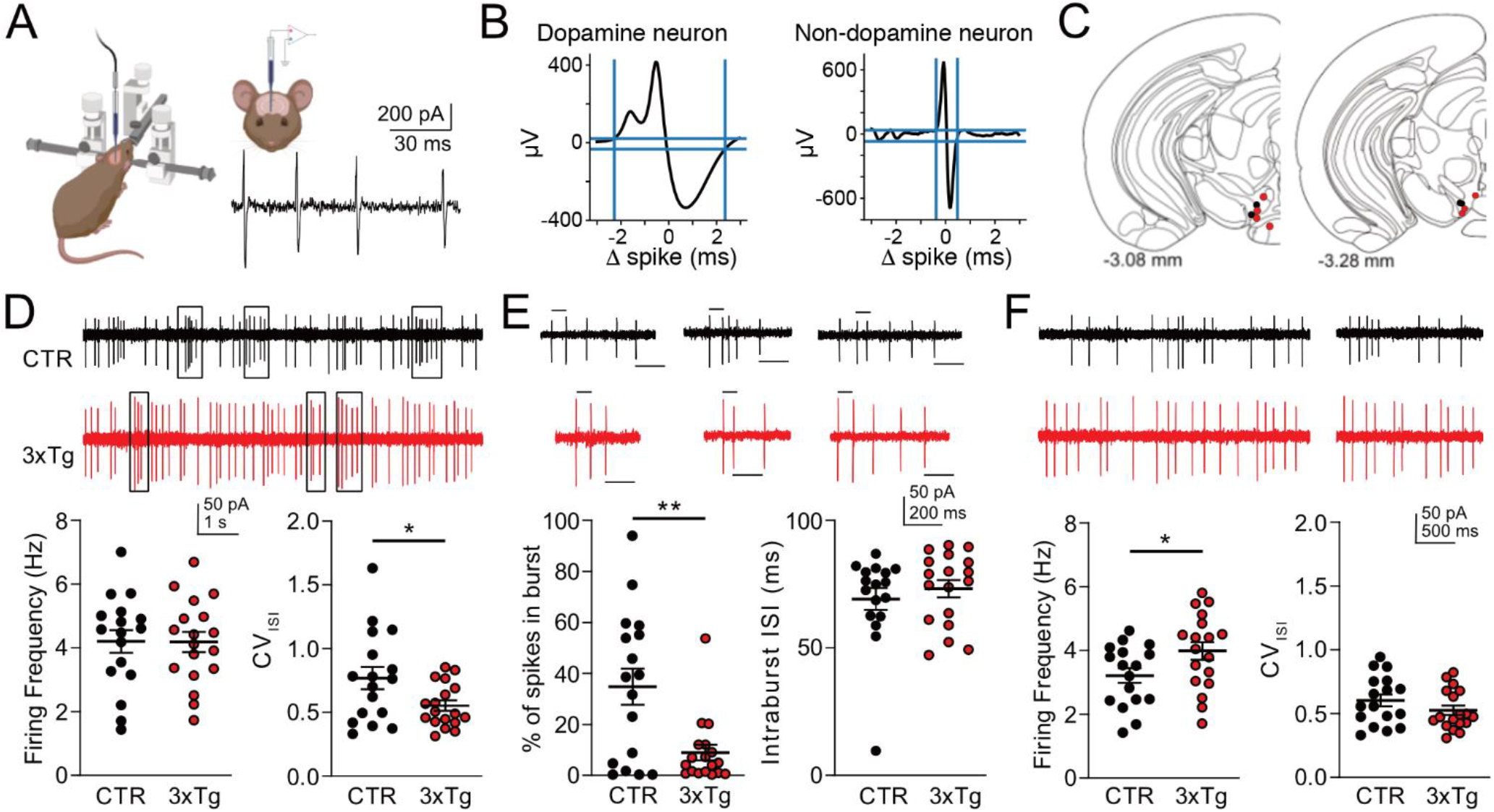
Impaired dopamine neuron bursting *in vivo* in 3xTg-AD mice. **A**. Diagram of in-vivo recordings of anesthetized mice. **B**. Sample dopamine neuron trace identification and rejection. **C**. Pontamine dot locations on final tracks in control (black) and 3xTg-AD (red) animals. **D**. Control (CTR, black) and 3xTg-AD (red) dopamine neurons fired at a similar rate (bottom left), but had a higher CV_ISI_ (bottom right). **E**. Bursts (from rectangles in D) that initiate with a first ISI < 80 ms (bar above) and a final ISI > 160 ms (bar below). The total number of spikes in bursts were significantly greater in 3xTg-AD dopamine neurons (bottom left), but fired at the same rate within a burst (bottom right). **F**. Spikes not in bursts (from D) showed that 3xTg-AD dopamine neurons fire faster than CTR mice (left) but show similar CV_ISI_s (right). CTR n = 16, 3xTg n = 18. Graphs show mean ± SEM; ^*^ = P < 0.05; ^**^ = P < 0.01.

We previously observed reward learning deficits in 3xTg-AD mice [5], a behavior that is highly reliant on dopamine neuron bursting and subsequent phasic release [11-15]. We therefore wondered if burst properties were altered in 3xTg-AD mice. We identified whether a spike was in a burst based on the commonly applied 80/160 ms criteria (see 2.2; [12]). Based on this, 3xTg-AD dopamine neurons showed a lower proportion of spikes within bursts (3xTg: 9 ± 3%; WT: 35 ± 7%), but similar inter-spike intervals within the bursts (3xTg: 70 ± 3 ms; WT 70 ± 4 ms; **Fig. 1E**). In contrast, in the fraction of spikes classified as “non-burst” (i.e. inter-burst), 3xTg-AD dopamine neurons showed an increase in firing frequency (3xTg: 4.0 ± 0.3 Hz; WT: 3.2 ± 0.2 Hz), with no difference in the CV_ISI_(3xTg: 0.53 ± 0.04; WT 0.60 ± 0.05; **Fig. 1F**). The difference in non-burst firing is consistent with data we previously obtained in dopamine neurons in brain slices, which do not burst spontaneously partially due to the relative lack of burst-driving synaptic input in slice preparations [5]. Thus, our *in vivo* data indicate that 3xTg-AD VTA dopamine neurons show a decrease in burst firing, but fire faster when not in a defined burst.

### 3.2 Impaired bursting of 3xTg-AD dopamine neurons is preserved *ex vivo*

The hyperexcitability of dopamine neurons in AD models is accompanied by a reduction in dopamine release and a decrease in reward learning [4, 5, 29]. This is in contrast to reports of ion channel manipulations that increased both burst firing and elevated firing rates, ultimately leading to more dopamine release [30]. *In vivo*, we observed fewer spikes within bursts in 3xTg-AD mice, yet neurons fired faster outside of defined bursts (**Fig. 1**). To explain this apparent paradox, we posited that single dopamine neurons from 3xTg-AD mice may be impaired in their ability to sustain burst firing. To test this, we prepared brain slices from 3xTg-AD and wild type mice and used depolarizing current steps to mimic bursting activity [29]. We previously showed that 3xTg-AD dopamine neurons have increased sensitivity to injected current [5], but those results could have been affected by dialysis of the cytoplasm using the whole-cell configuration. To more accurately assess bursting during sustained depolarization, we used the minimally-invasive gramicidin perforated patch technique to record medial VTA dopamine neurons, and injected 2-second current steps to assess action potential generation in response to sustained excitation. While there are several experimental paradigms that could simulate bursting, this is among the clearest paradigm to evaluate adaptation, maximum firing frequency, and sensitivity to current [16, 18]. To measure the overall sensitivity to injected current, we applied 25 pA steps from -100 to +200 pA and calculated the firing response (**Fig. 2A**). We found 3xTg-AD dopamine neurons exhibit an increased sensitivity to small current steps, however, this was lost at injection amplitudes in excess of +100 pA. We also observed that at higher current injections, 3xTg-AD neurons were more likely to cease firing within a second of depolarization (**Fig. 2A,B**). To further investigate this, we divided the depolarization into two one-second halves (S1 and S2) and assessed the firing rates separately. In the first second of depolarization, 3xTg-AD dopamine neurons showed hyperexcitability (**Fig. 2B)**, as we have described [5]. When we compared S2 to S1, we saw minimal change in wild type mice, but a profound decrease of action potential number in 3xTg-AD neurons beginning with the +100 pA step (**Fig. 2C,D**). These data suggest that depolarized 3xTg-AD dopamine neurons initially fire faster than wild type neurons but stop firing sooner, preventing sustained firing of action potentials within the simulated burst.

**Figure 2.**
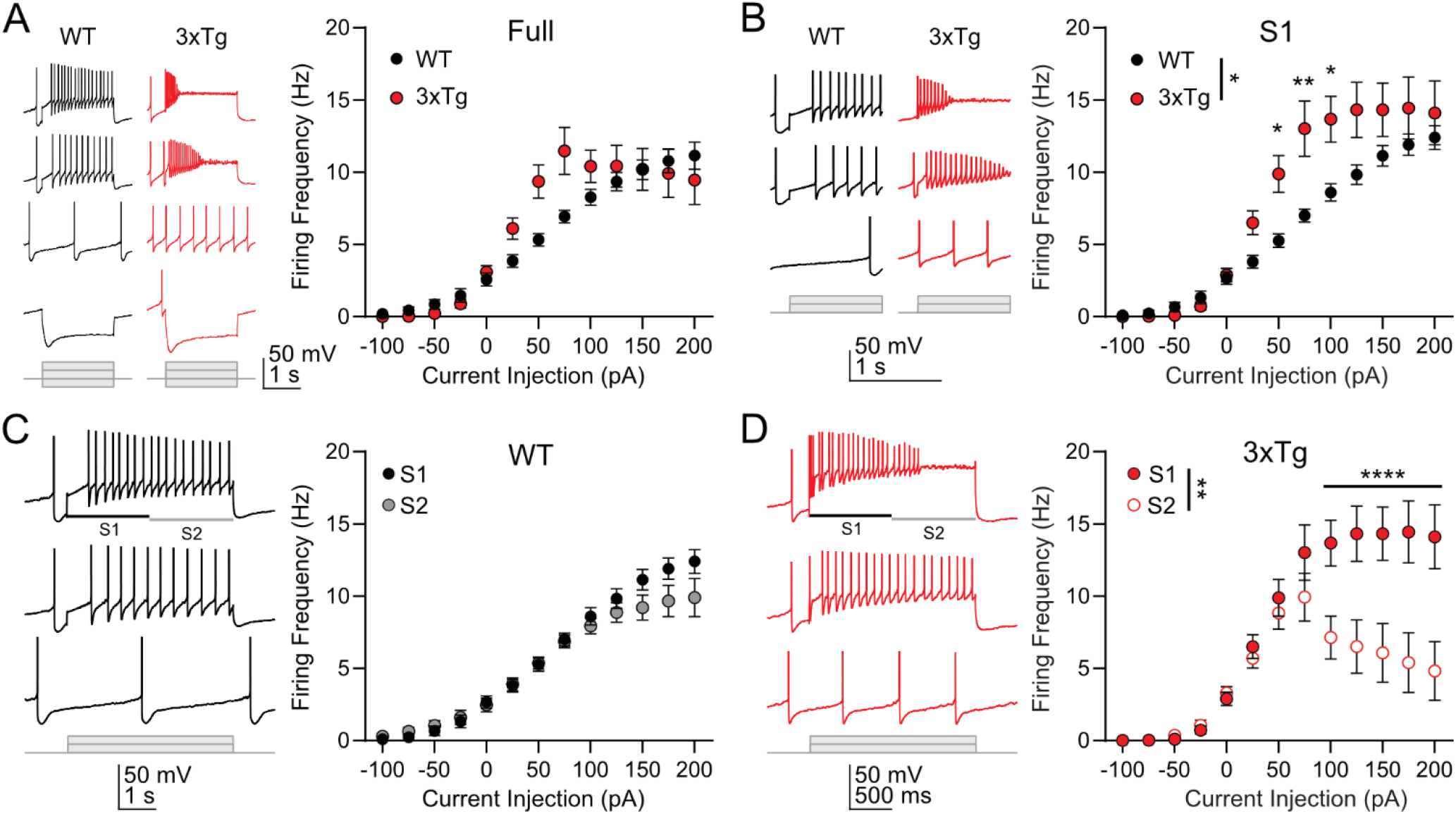
Impaired VTA dopamine neuron firing in 3xTg-AD brain slices. **A**. Frequency-current (F-I) plot showing the number of spikes per second given each current injection step from -100 pA to + 200 pA. There was a significant interaction between the current and firing frequency based on the genotype (P = 0.002), but there was no main effect of genotype between the WT and 3xTg-AD neurons (2-Way RM ANOVA genotype F_1,19_= 1; P = 0.3). **B**. 3xTg-AD F-I curves showed an increase in sensitivity to current during the first second of the current step (2-Way RM ANOVA genotype F_1,19_ = 4.6, P = 0.04; genotype x current P < 0.0001). **C**. WT neurons showed slight adaptation in S2 of the current step with high current steps showing different response to current (current x time interaction; P < 0.0001) but no main effect between S1 and S2 (2-Way RM ANOVA S F_1,9_ = 3.6, P = 0.09). **D**. 3xTg-AD neurons exhibited decreased firing in S2 at current injections ≥ 100 pA (2-Way RM ANOVA F_1,10_ = 12, P = 0.006; current x time interaction; P < 0.0001). WT n = 10, 3xTg-AD n = 11. Mean ± SEM;^*^ = P < 0.05; ^**^ = P < 0.01, ^****^ = P < 0.0001.

### 3.3 Sustained depolarization impairs bursting

Depolarization block is a well described phenomenon that limits the average firing frequency of dopamine neurons to approximately 10 Hz at steady state [17]. The slow inactivation of active voltage-gated sodium channels occurs at membrane potentials near action potential threshold with a time constant in the tens to hundreds of milliseconds and recovers slowly [17, 31]. Since prolonged depolarization increases the likelihood of inactivation of voltage-gated sodium channels [16, 17], we tested whether sustained depolarization affects the ability of dopamine neurons to sustain bursting. We obtained a stable perforation on wild type VTA dopamine neurons and determined the baseline sensitivity to current using an F-I curve as in Figure 2. We next injected a positive holding current (+25 to +125 pA) to reversibly depolarize the minimum voltage trajectory by about +10 mV (baseline: -74 ± 2 mV, depolarization: -62 ± 3 mV, post: -73 ± 4 mV; **Fig. 3A**), similar to what we previously observed in 3xTg-AD mice [5]. The sustained positive current injection depolarized the action potential threshold (baseline: -56.4 ± 2.3 mV, positive current: -51.5 ± 3.1 mV; **Fig. 3B**) suggesting sodium channel inactivation was likely present. Positive current injection also produced faster firing (baseline: 2.2 ± 0.4 Hz, depolarization: 5.2 ± 0.8 Hz) but without disrupting the fidelity of pacemaking (CV_ISI_ baseline: 0.10± 0.04, depolarization: 0.11 ± 0.03; **Fig. 3B**). When assessing bursting with F-I curves, depolarization block was not observed with small current injections (+50 pA or less), but the depolarized neurons ceased firing during large positive current injections (**Fig. 3C**). Firing parameters returned to baseline levels after the depolarizing current was released. These results obtained in wild type mice support the notion that the baseline depolarized state observed in 3xTg-AD dopamine neurons may limit burst firing through depolarization block.

**Figure 3.**
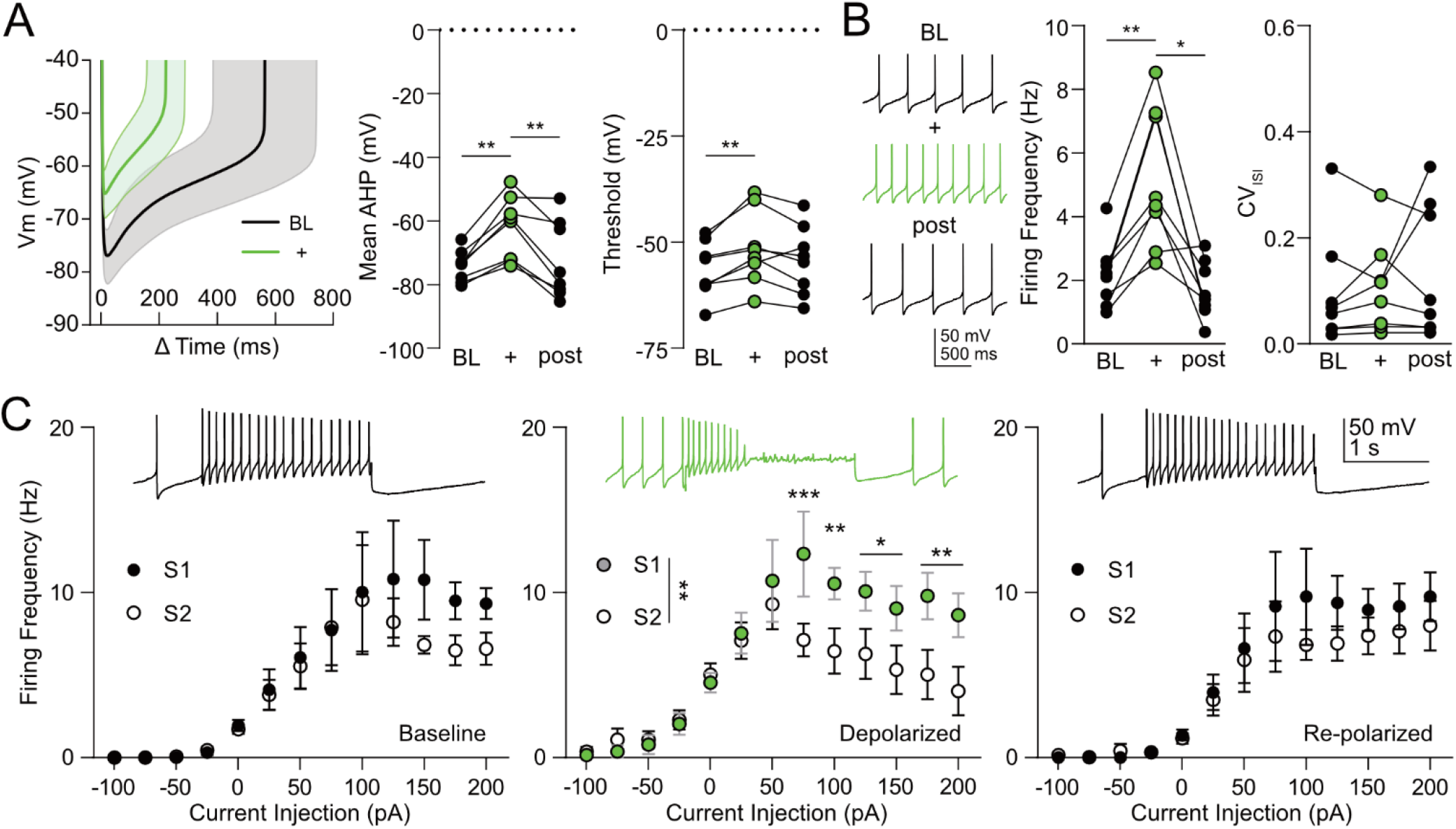
Sustained depolarization increases adaptation in VTA dopamine neurons. **A**. Injecting positive holding current into dopamine neurons from wild type mice (+, green) depolarized their mean AHP voltage and was reversible (middle; RM 1-Way ANOVA F_1.5,10_ = 13, P = 0.003). Action potential thresholds were also slightly depolarized (right; RM 1-Way ANOVA F_1.8,13_ = 8, P = 0.007) **B**. Firing frequency from depolarized cells increased firing frequency (middle; RM 1-Way ANOVA F_1.2,8_ = 18, P = 0.002) but did not affect the CV_ISI_ (right; RM 1-Way ANOVA F_1.3,9_ = 0.8, P = 0.4). **C**. F-I curves during the baseline phase showed slight adaptation late in the current step (left; 2-Way RM ANOVA second F_1,7_ = 3, P = 0.1; time x current interaction P = 0.008), but sustained depolarization produced a stronger difference (middle; 2-Way RM ANOVA second F_1,7_ = 18, P = 0.004), which returned to baseline (right; 2-Way RM ANOVA second F_1,7_ = 3, P = 0.1; time x current interaction P = 0.05). Biological N = 5; n = 8. Mean ± SEM; ^*^ = P < 0.05, ^**^ = P < 0.01, ^***^ = P <0.001.

### 3.4 Aβ does not contribute to dopamine neuron firing aberrations

3xTg-AD mice express both amyloid and tau pathology. While we previously and presently observed clear differences in dopamine neuron activity between WT and 3xTg-AD mice [5], we wondered whether Aβ or tau was predominantly responsible for the aberrations observed in intrinsic and burst firing. To isolate the effects of Aβ accumulation in absence of hyperphosphorylated hTau, we employed the APP^NL-G-F^ knock-in model of amyloid beta under the endogenous amyloid precursor protein (APP) promoter [19]. We first examined the accumulation of Aβ plaques by staining with methoxy-X04. As expected, we observed extracellular Aβ plaques throughout the cortex and hippocampus in 6-month-old APP^NL-G-F^ mice. Remarkably, we also observed robust accumulation of amyloid within the VTA of APP^NL-G-F^, but not 12-month-old 3xTg-AD mice (**Fig. 4A**). Overall, the plaque accumulation in the APP^NL-G-F^ line was pervasive throughout the brain, as previously reported [32].

**Figure 4.**
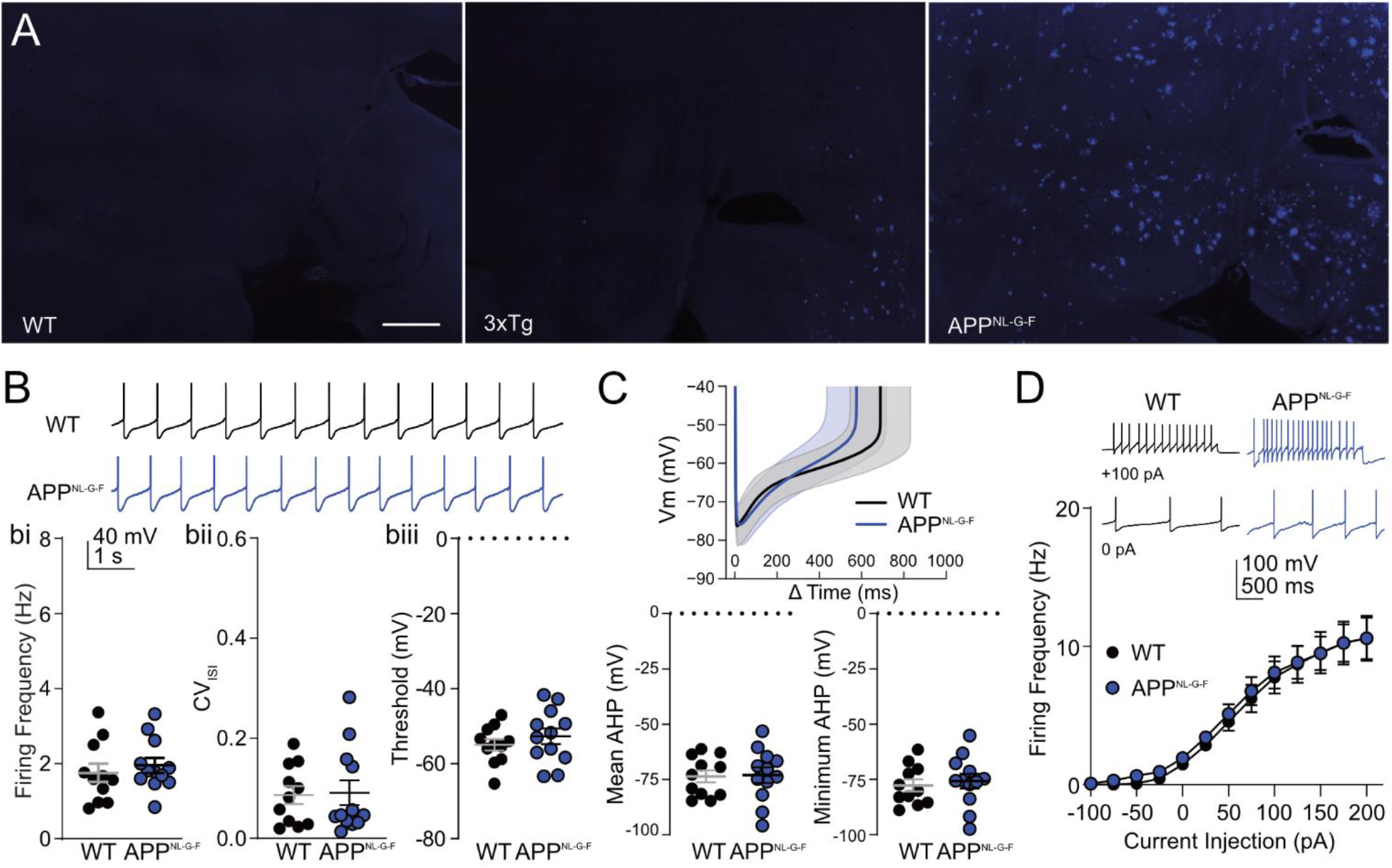
APP^NL-G-F^ VTA dopamine neurons do not exhibit increased firing or depolarization. **A**. Methoxy-X04 staining revealed increased VTA amyloid beta plaque burden in 6-month-old APP^NL-G-F^ mice versus 3xTg-AD mice and WT controls. Scale bar = 500 µm **B**. Intrinsic firing properties of wildtype C57Bl/6 (WT; black) and APP^NL-G-F^ mice (blue). **bi** firing frequency was not different between genotypes (t_21_ = 0.6, P = 0.5). **bii** CV_ISI_ was not different (Mann-Whitney U = 65, P ≈ 1). **Biii** AP threshold was also not different (t_21_, = 0.9, P = 0.4). **C**. Voltage trajectories were not different between WT and APP^NL-G-F^ dopamine neurons either measured by mean AHP (left, t_21_ = 0.1, P = 0.9) or minimum ramp voltage (middle, t_21_ = 0.5, P = 0.7). **D**. F-I curves showed no differences in APP^NL-G-F^ dopamine neuron sensitivity to current (2-way RM ANOVA genotype F_1,20_ = 0.1, P = 0.7; genotype x current P ≈ 1). WT n = 11-12 APP^NL-G-F^ n = 11-12. Mean ± SEM

We next investigated dopamine neuron intrinsic firing in APP^NL-G-F^ mice, which we previously showed is disrupted in the 3xTg-AD model [5]. We recorded from dopamine cells using gramicidin perforated patch and found no change in firing frequency from control mice (APP 2.0 ± 0.2 Hz, WT 1.8± 0.2 Hz; **Fig. 4Bi**). We also observed no difference in the regularity of the firing (CV_ISI_; APP 0.09 ± 0.03, WT 0.09 ± 0.02; **Fig. 4bii**), or the action potential threshold (APP -53 ± 2 mV, WT -55 ± 2 mV; **Fig. 4biii**). Additionally, there was no difference in the inter-spike trajectory (**Fig. 4C**), including the average afterhyperpolarization potential (AHP; APP mice -73.0 ± 3.5 mV, WT -73.7 ± 2.7 mV), or the minimum AHP (APP -75.7 ± 3.3 mV, WT -77.7 ± 2.7 mV). We also observed no difference in sensitivity to injected current between WT and APP^NL-G-F^ mice (**Fig. 4D**). These data indicate that despite clear accumulation of plaques within the VTA, dopamine neurons in APP^NL-G-F^ mice do not differ in their intrinsic pacemaking or in their capacity to burst.

### 3.5 hTau expression alters dopamine neuron firing

Consistent with human AD, 3xTg-AD mice also exhibit excessive accumulation of intracellular neurofibrillary (tau) tangles throughout the brain [20]. We therefore sought to determine the effects of hyperphosphorylated human tau specifically expressed in dopamine neurons. To do this, we stereotaxically injected a Cre-dependent virus to express hTau (AAV-DIO-hTau_P364S_) with a mutation that renders it prone to hyperphosphorylation. Injections were delivered into the VTA of mice expressing Cre recombinase under the dopamine transporter promoter locus (DAT^Cre^). AAV5-DIO-eYFP injection into DAT^Cre^ mice was used as a control. We confirmed phosphorylated tau (pTau) expression in VTA dopamine cell bodies of AAV-hTau injected mice by immunofluorescence (**Fig.5A**). We then recorded the spontaneous firing of hTau-transduced VTA dopamine neurons ∼4 weeks after surgery. Similar to what we previously observed in 3xTg-AD mice, hTau expression increased the firing rate of VTA dopamine neurons (hTau 3.1 ± 0.3 Hz, CTR 2.1 ± 0.3 Hz; **Fig. 5bi**) and produced more irregular firing (CV_ISI_: hTau 0.13 ± 0.03, CTR 0.05 ± 0.004; **Fig. 5bii**). Also similar to the 3xTg-AD model, we saw depolarization of the action potential threshold (hTau -46.7 ± 2.4 mV, CTR -55.0 ± 1.8 mV; **Fig. 5biii**), indicating possible increased inactivation of voltage-gated sodium channels. To further investigate the alterations in pacemaking, we analyzed the inter-spike voltage trajectories (**Fig. 5C**). hTau expression depolarized the inter-spike trajectory, including the average AHP (Tau -59.3 ± 3.3 mV, CTR -72.3 ± 2.3 mV) and the minimum voltage trajectory (Tau -64.0 ± 3.4 mV, CTR -76.8 ± 2.4 mV, **Fig. 5C**). Thus, introduction of intracellular hTau increases firing frequency and induces more irregular firing, similar to what we previously observed in 3xTg-AD mice [5].

**Figure 5.**
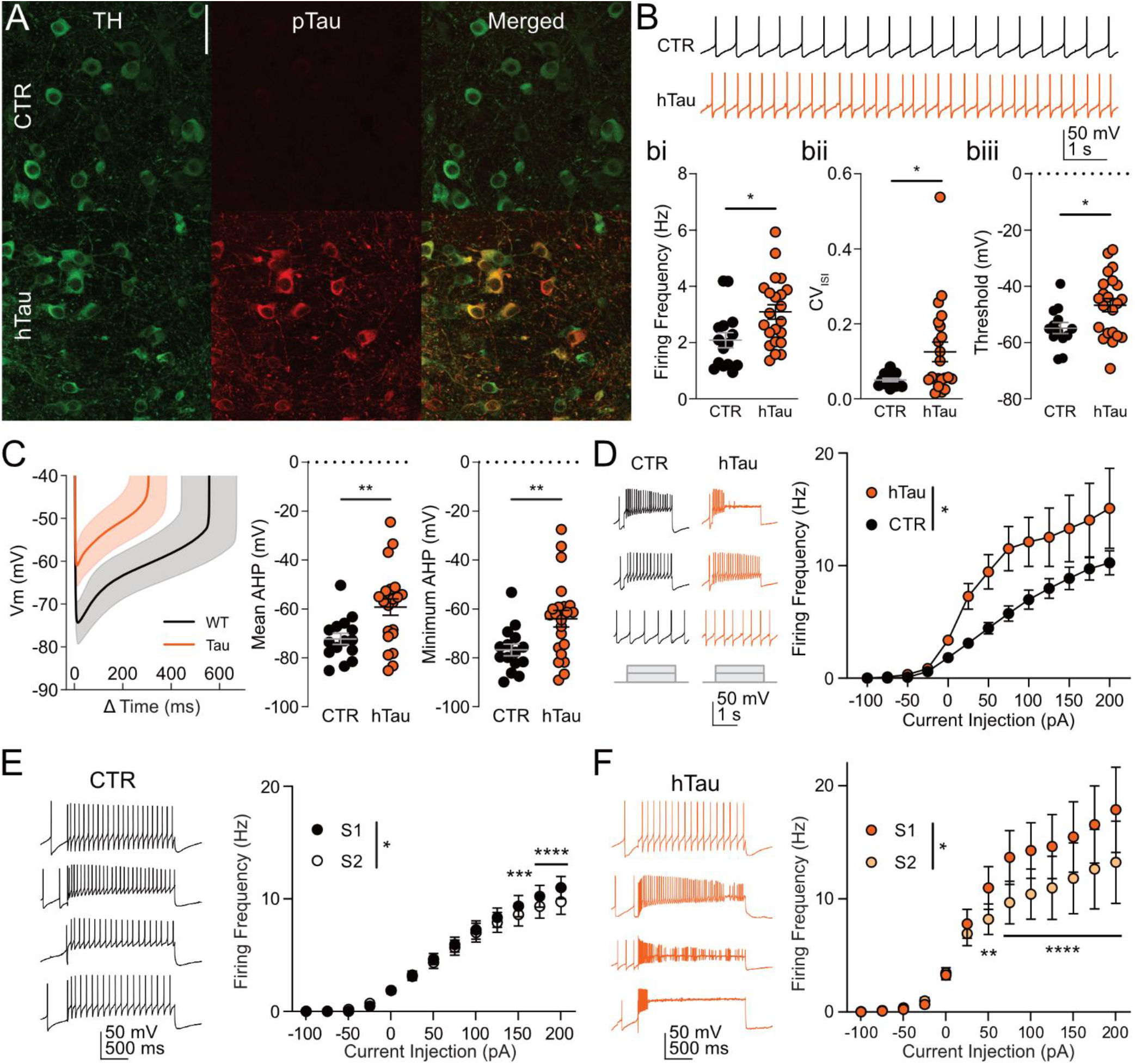
The presence of phosphorylated hTau alters VTA dopamine neuron firing. **A**. Phosphorylated tau was induced after stereotaxic injection of AAV-DIO-hTau_P364S_ (scale bar = 50 µm). **B**. Intrinsic firing properties of control DAT^Cre^ or DAT^Cre^-EYFP mice (CTR; black) and hTau mice (red). **bi** Firing of dopamine neurons in hTau mice became faster with hTau (t_35_ = 2.6, P = 0.01). **bii**. Firing in the presence of hTau was more irregular (U = 101, P = 0.049). **biii**. Threshold of action potentials was depolarized in hTau mice (t_35_ = 2.6, 0.2). **C**. The ramp of the inter-spike interval (left) was depolarized when either looking at the mean AHP (t_35_ = 2.9, P = 0.006) or minimum AHP (t_35_ = 2.8, P = 0.009). **D**. F-I curves showed an increase of sensitivity to current injection in hTau mice (2-way RM ANOVA F_1,31_ = 4, P = 0.049). **E**. Left, firing in four example dopamine neurons at +200 pA. Control mice showed a slight adaptation at high current injections +150 pA (2-way RM ANOVA second F_1,14_ = 5, P = 0.04; current x time interaction P < 0.0001). **F**. Left, firing in four example dopamine neurons at +200 pA. hTau mice experienced faster adaptation starting at +50 pA (2-way RM ANOVA second F_1,17_ = 11, P = 0.005; current x time interaction P < 0.0001). CTR n = 15, hTau n = 22). Mean ± SEM; ^*^ = P < 0.05, ^**^ = P < 0.01, ^***^ = P < 0.001, ^***^ = P < 0.0001.

Finally, we tested whether the bursting deficits we observed in 3xTg-AD mice (**Fig. 1, 2**) were also present in the hTau mice. We found that hTau expression did increase the sensitivity to injected current (**Fig. 5D**). Control dopamine neurons displayed a small adaptation at high depolarization steps (**Fig. 5E**). However, expression of hTau decreased the average firing rate in the second half of the depolarizing current steps at current injection amplitudes as small as +50 pA (**Fig. 5F**). During large current steps, we observed highly varied responses among individual neurons, ranging from normal firing to complete depolarization block (**Fig. 5F**) which may be explained in part due to variation of AAV transduction and/or hTau expression illustrated in **Fig. 5A**.

Taken together, our data show that the presence of hyperphosphorylated human tau, but not Aβ, recapitulates the dopamine neuron firing aberrations observed in the 3xTg-AD model. Further, dopamine neurons burdened with hTau are depolarized, inducing a hyperactive, faster firing state. This depolarization can partially limit the capability of the neuron to burst, possibly through inactivation of voltage-gated sodium channels. This impairment of bursting seen in *ex vivo* slice recordings might also explain the smaller proportion of spikes in bursts observed *in vivo*.

## 4 DISCUSSION

Although investigations into dopamine neuron activity in AD mouse models are sparse, the published evidence suggests that dopamine neurons may be intrinsically hyperactive but exhibit decreased dopamine release [4-6]. This apparent paradox drew us to further investigate hTau- and amyloid-based AD models using *ex vivo* and *in vivo* paradigms. We first measured spontaneous activity of dopamine cells in anesthetized 3xTg-AD and wild type mice. We saw that 3xTg-AD dopamine neurons fire faster outside of classically defined bursts[12], but generate fewer burst spikes. Subsequent experiments conducted in brain slices indicated that 3xTg-AD dopamine neurons are hypersensitive to brief current injections as we previously reported [5], but more readily enter into depolarization block, thus limiting action potential generation during periods of sustained excitation.This result suggests that while intact neurons from 3xTg-AD mice may release more tonic dopamine, burst firing-mediated phasic release may be blunted. We further show that VTA dopamine neuron firing deficits are most likely caused by pathogenic tau expression, and that amyloid accumulation alone does not significantly alter intrinsic firing activity.

### 4.1 Mechanisms of firing alterations

Understanding the *in vivo* electrophysiological alterations produced in AD models may help to elucidate the neuronal underpinnings of behavioral phenotypes. We recently described deficits in VTA-dependent responding for food reward in the 3xTg-AD model [5]. While decreased dopamine release could be caused by a decrease in dopamine neuron or terminal number [4], we posit this could also result from an inability to sustain bursts of action potentials to induce phasic dopamine release (see **Fig. 3**). Future experiments investigating behavior-dependent release will be required to fully parse our *in vivo* and *ex vivo* observations. We note that while sustained depolarization in wild type mice (**Fig. 3**) effectively mimics the adaptation observed in 3xTg-AD mice, it does *not* increase the sensitivity to current, based on the unchanged firing rate in response to small current injections of the F-I curve. Further, while sustained depolarization increases firing rate, it does not alter CV_ISI_. So, while our data indicate that depolarization contributes to enhanced depolarization block in the 3xTg-AD model [16, 17], other molecular alterations are likely responsible for additional changes in firing. We previously reported the contribution of enhanced casein kinase 2 activity and decreased SK channel currents in VTA dopamine neurons in 3xTg-AD mice. When SK channels were blocked, dopamine neurons also depolarized to a similar extent [5]. Therefore, our current results extend these findings by showing that neuronal depolarization produced by sustained current injection only partially replicates the fast, irregular firing observed in brain slices, yet ultimately explains the firing patterns of 3xTg-AD mice *in vivo*.

### 4.2 Amyloid β and dopamine neuron firing

In APP^NL-G-F^ VTA dopamine neurons, we saw no disruption of intrinsic firing. Conversely, studies in the hippocampus and cortex highlight changes in neuronal firing in single-neuron and population recordings [33, 34]. Experiments from parvalbumin interneurons in the lateral entorhinal cortex show similar firing disruptions in response to depolarizing current steps [33]. Although other models of AD exhibit minimal Aβ plaque formation near the VTA, 6-month-old APP^NL-G-F^ mice clearly display Aβ plaques. Therefore, we can likely rule out the idea that the absence of firing deficits is due to a lack of Aβ plaques in the immediate area. In conjunction with intrinsic deficits previously observed in the APP^NL-G-F^ mice, bidirectional synaptic changes have been reported in the hippocampus, entorhinal cortex, and prefrontal cortex [33, 35-37]. Thus, while there is little evidence of impaired dopamine neuron firing, synaptic changes could be present that we have not yet investigated.Regardless, our results suggest that Aβ alone is insufficient to induce dopamine neuron firing deficits.

### 4.3 hTau and dopamine neuron firing

To eliminate nonspecific effects of hTau accumulation, we delivered a Cre-dependent viral vector to transduce dopamine neurons specifically with hTau. While a limitation of this model is the potential for varying levels of hTau expression, we observed prominent firing deficits despite this heterogeneity (**Fig. 5**). In contrast to our findings, in CA1 pyramidal neurons, hTau expression decreases firing rates across a range of injected currents, even though the action potential thresholds are depolarized, similar to our findings [38]. CA1 pyramidal neurons do not intrinsically pacemake like dopamine neurons and therefore may respond to the added molecular stress of hTau in wholly different manner. For example, that study reported that adaptation to injected current occurs in cultured primary hippocampal neurons [38], similar to our findings. The decreased excitability of CA1 neurons in AD models is accompanied by widely reported deficits in synaptic transmission and plasticity [39, 40]. Dopaminergic dysfunction may be a secondary contributor to synaptic deficits elsewhere in the brain, as dopamine release from the axon terminals of VTA neurons regulates synaptic plasticity, learning, and memory in several brain regions [4, 11, 41-43].

Future studies are required to identify the mechanisms of altered dopamine neuron firing caused by tau pathology and downstream signaling effects. While our previous work attributed disrupted VTA dopamine neuron firing to CK2 dysregulation of SK channels [5], more work is needed to uncover the pathway by which hTau may increase CK2 activity at the SK channel complex. Similarly, dopaminergic signaling is directly associated with formation and maintenance of long-term memory [4, 11, 30, 41, 43]. This memory, including that induced by reward learning, is known to require bursting of dopamine neurons [11-15, 30]. Therefore, if bursting of VTA dopamine neurons is affected through pathological accumulation of tau, impairment of learning may precede neuronal degeneration and death in projection areas.

### 4.4 Amyloid β – hTau interactions

Although we have identified hTau as the mediator of intrinsic firing aberrations in VTA dopamine neurons, Aβ and phosphorylated tau likely interact to produce the neuronal and circuit alterations responsible for AD symptoms. One general hypothesis is that Aβ and phosphorylated tau accelerate each other’s pathology. These interactions ultimately cause irreversible damage to neurons or circuits through mitochondrial, synaptic, or inflammatory actions [44]. Both *in vivo* and *ex vivo* studies of AD models suggest that while hyperexcitability is produced by Aβ synaptopathies, the effects can be overwhelmed by the strong neuronal activity suppression induced by hyperphosphorylated hTau [45, 46]. Given that action potentials are the output of the neuron, there is a case that tau impairs dopamine neurons strongly regardless of potential Aβ deficits in synaptic regulation. Interestingly, our findings suggest a two-pronged tauopathy in the VTA that may bidirectionally affect neuronal output. Our results highlight that hyperphosphorylated hTau, but not Aβ, mediates hyperexcitability of VTA dopamine neurons but also enhances action potential cessation through depolarization block.

## AUTHOR CONTRIBUTION

Conceptualization and Methodology: W.M.K., E.T-R. and M.J.B; Formal Analysis and Investigation: W.M.K., M.H.H.; Writing-Original Draft: W.M.K. and E.T-R.; Contribution of Research Materials: D.W. and H.C.R.; Writing-Review and Editing: W.M.K., E.T-R., H.E.B., M.H.H., A.K., D.W. H.C.R., and M.J.B; Supervision and Project Administration: M.J.B.

## ACKNOWLEDGEMENTS

We thank the Ocañas lab for their help with MX-04 staining protocols. We thank all members of the Beckstead lab for helpful comments throughout the project, especially Kelsey Carter and Nicole Yates for technical assistance. This work was supported by NIH R01 NS135830 (MJB), RF1 AG079199 (DW), F31 AG079620 (HEB) and Veterans Affairs Merit grant I01BX005396 (MJB). WMK was supported by NIH training grant T32 AG052363.

## REFERENCES

[1] Botto R, Callai N, Cermelli A, Causarano L, Rainero I. Anxiety and depression in Alzheimer’s disease: a systematic review of pathogenetic mechanisms and relation to cognitive decline. Neurol Sci. 2022;43:4107–24.

[2] Colella D, Guerra A, Paparella G, Cioffi E, Di Vita A, Trebbastoni A, et al. Motor dysfunction in mild cognitive impairment as tested by kinematic analysis and transcranial magnetic stimulation. Clin Neurophysiol. 2021;132:315–22.

[3] Phan SV, Osae S, Morgan JC, Inyang M, Fagan SC. Neuropsychiatric Symptoms in Dementia: Considerations for Pharmacotherapy in the USA. Drugs in R&D. 2019;19:93–115.

[4] Nobili A, Latagliata EC, Viscomi MT, Cavallucci V, Cutuli D, Giacovazzo G, et al. Dopamine neuronal loss contributes to memory and reward dysfunction in a model of Alzheimer’s disease. Nat Commun. 2017;8:14727.

[5] Blankenship HE, Carter KA, Pham KD, Cassidy NT, Markiewicz AN, Thellmann MI, et al. VTA dopamine neurons are hyperexcitable in 3xTg-AD mice due to casein kinase 2-dependent SK channel dysfunction. Nature Communications. 2024;15.

[6] Krashia P, Nobili A, D’Amelio M. Unifying Hypothesis of Dopamine Neuron Loss in Neurodegenerative Diseases: Focusing on Alzheimer’s Disease. Front Mol Neurosci. 2019;12:123.

[7] Krashia P, Spoleti E, D’Amelio M. The VTA dopaminergic system as diagnostic and therapeutical target for Alzheimer’s disease. Front Psychiatry. 2022;13:1039725.

[8] Volman SF, Lammel S, Margolis EB, Kim Y, Richard JM, Roitman MF, et al. New insights into the specificity and plasticity of reward and aversion encoding in the mesolimbic system. J Neurosci. 2013;33:17569–76.

[9] Delgado MR, Dickerson KC. Reward-related learning via multiple memory systems. Biol Psychiatry. 2012;72:134–41.

[10] Qi G, Zhang P, Li T, Li M, Zhang Q, He F, et al. NAc-VTA circuit underlies emotional stress-induced anxiety-like behavior in the three-chamber vicarious social defeat stress mouse model. Nat Commun. 2022;13:577.

[11] Speranza L, di Porzio U, Viggiano D, de Donato A, Volpicelli F. Dopamine: The Neuromodulator of Long-Term Synaptic Plasticity, Reward and Movement Control. Cells. 2021;10:735.

[12] Grace AA, Bunney BS. Intracellular and extracellular electrophysiology of nigral dopaminergic neurons--1. Identification and characterization. Neuroscience. 1983;10:301–15.

[13] Aberman JE, Ward SJ, Salamone JD. Effects of Dopamine Antagonists and Accumbens Dopamine Depletions on Time-Constrained Progressive-Ratio Performance. Pharmacology Biochemistry and Behavior. 1998;61:341–8.

[14] Sokolowski JD, Salamone JD. The Role of Accumbens Dopamine in Lever Pressing and Response Allocation: Effects of 6-OHDA Injected into Core and Dorsomedial Shell. Pharmacology Biochemistry and Behavior. 1998;59:557–66.

[15] Nowend KL, Arizzi M, Carlson BB, Salamone JD. D1 or D2 antagonism in nucleus accumbens core or dorsomedial shell suppresses lever pressing for food but leads to compensatory increases in chow consumption. Pharmacology Biochemistry and Behavior. 2001;69:373–82.

[16] Blythe SN, Wokosin D, Atherton JF, Bevan MD. Cellular mechanisms underlying burst firing in substantia nigra dopamine neurons. J Neurosci. 2009;29:15531–41.

[17] Qian K, Yu N, Tucker KR, Levitan ES, Canavier CC. Mathematical analysis of depolarization block mediated by slow inactivation of fast sodium channels in midbrain dopamine neurons. J Neurophysiol. 2014;112:2779–90.

[18] Grace AA, Bunney BS. The control of firing pattern in nigral dopamine neurons: burst firing. J Neurosci. 1984;4:2877–90.

[19] Saito T, Matsuba Y, Mihira N, Takano J, Nilsson P, Itohara S, et al. Single App knock-in mouse models of Alzheimer’s disease. Nat Neurosci. 2014;17:661–3.

[20] Oddo S, Caccamo A, Shepherd JD, Murphy MP, Golde TE, Kayed R, et al. Triple-transgenic model of Alzheimer’s disease with plaques and tangles: intracellular Abeta and synaptic dysfunction. Neuron. 2003;39:409–21.

[21] Dominguez-Lopez S, Sharma R, Beckstead MJ. Neurotensin receptor 1 deletion decreases methamphetamine self-administration and the associated reduction in dopamine cell firing. Addiction Biology. 2019;26.

[22] Paxinos GaF, K.B.J.. The Mouse Brain in Stereotaxic Coordinates. 2nd Edition ed. San Diego: Academic Press; 2001.

[23] Grace AA, Bunney BS. Intracellular and extracellular electrophysiology of nigral dopaminergic neurons--3. Evidence for electrotonic coupling. Neuroscience. 1983;10:333–48.

[24] Dominguez-Lopez S, Ahn B, Sataranatarajan K, Ranjit R, Premkumar P, Van Remmen H, et al. Long-term methamphetamine self-administration increases mesolimbic mitochondrial oxygen consumption and decreases striatal glutathione. Neuropharmacology. 2023;227:109436.

[25] Khaliq ZM, Bean BP. Pacemaking in dopaminergic ventral tegmental area neurons: depolarizing drive from background and voltage-dependent sodium conductances. J Neurosci. 2010;30:7401–13.

[26] Tarfa RA, Evans RC, Khaliq ZM. Enhanced Sensitivity to Hyperpolarizing Inhibition in Mesoaccumbal Relative to Nigrostriatal Dopamine Neuron Subpopulations. J Neurosci. 2017;37:3311–30.

[27] Gantz SC, Ford CP, Morikawa H, Williams JT. The Evolving Understanding of Dopamine Neurons in the Substantia Nigra and Ventral Tegmental Area. Annu Rev Physiol. 2018;80:219–41.

[28] Marinelli M, McCutcheon JE. Heterogeneity of dopamine neuron activity across traits and states. Neuroscience. 2014;282:176–97.

[29] La Barbera L, Nobili A, Cauzzi E, Paoletti I, Federici M, Saba L, et al. Upregulation of Ca(2+)-binding proteins contributes to VTA dopamine neuron survival in the early phases of Alzheimer’s disease in Tg2576 mice. Mol Neurodegener. 2022;17:76.

[30] Juarez B, Kong M-S, Jo YS, Elum JE, Yee JX, Ng-Evans S, et al. Temporal scaling of dopamine neuron firing and dopamine release by distinct ion channels shape behavior. Science Advances. 2023;9:eadg8869.

[31] Knowlton CJ, Ziouziou TI, Hammer N, Roeper J, Canavier CC. Inactivation mode of sodium channels defines the different maximal firing rates of conventional versus atypical midbrain dopamine neurons. PLoS Comput Biol. 2021;17:e1009371.

[32] Whyte LS, Hemsley KM, Lau AA, Hassiotis S, Saito T, Saido TC, et al. Reduction in open field activity in the absence of memory deficits in the App(NL-G-F) knock-in mouse model of Alzheimer’s disease. Behav Brain Res. 2018;336:177–81.

[33] Petrache AL, Rajulawalla A, Shi A, Wetzel A, Saito T, Saido TC, et al. Aberrant Excitatory-Inhibitory Synaptic Mechanisms in Entorhinal Cortex Microcircuits During the Pathogenesis of Alzheimer’s Disease. Cereb Cortex. 2019;29:1834–50.

[34] Arroyo-Garcia LE, Isla AG, Andrade-Talavera Y, Balleza-Tapia H, Loera-Valencia R, Alvarez-Jimenez L, et al. Impaired spike-gamma coupling of area CA3 fast-spiking interneurons as the earliest functional impairment in the App(NL-G-F) mouse model of Alzheimer’s disease. Mol Psychiatry. 2021;26:5557–67.

[35] Latif-Hernandez A, Sabanov V, Ahmed T, Craessaerts K, Saito T, Saido T, et al. The two faces of synaptic failure in App(NL-G-F) knock-in mice. Alzheimers Res Ther. 2020;12:100.

[36] Carpanini SM, Torvell M, Bevan RJ, Byrne RAJ, Daskoulidou N, Saito T, et al. Terminal complement pathway activation drives synaptic loss in Alzheimer’s disease models. Acta Neuropathol Commun. 2022;10:99.

[37] Blume T, Filser S, Sgobio C, Peters F, Neumann U, Shimshek D, et al. beta-secretase inhibition prevents structural spine plasticity deficits in App (NL-G-F) mice. Front Aging Neurosci. 2022;14:909586.

[38] Hatch RJ, Wei Y, Xia D, Gotz J. Hyperphosphorylated tau causes reduced hippocampal CA1 excitability by relocating the axon initial segment. Acta Neuropathol. 2017;133:717–30.

[39] Yoshiyama Y, Higuchi M, Zhang B, Huang SM, Iwata N, Saido TC, et al. Synapse loss and microglial activation precede tangles in a P301S tauopathy mouse model. Neuron. 2007;53:337–51.

[40] Wu M, Zhang M, Yin X, Chen K, Hu Z, Zhou Q, et al. The role of pathological tau in synaptic dysfunction in Alzheimer’s diseases. Transl Neurodegener. 2021;10:45.

[41] Rossato JI, Bevilaqua LRM, Izquierdo I, Medina JH, Cammarota M. Dopamine Controls Persistence of Long-Term Memory Storage. Science. 2009;325:1017–20.

[42] Yagishita S, Hayashi-Takagi A, Ellis-Davies Gcr, Urakubo H, Ishii S, Kasai H. A critical time window for dopamine actions on the structural plasticity of dendritic spines. Science. 2014;345:1616–20.

[43] Broussard JI, Yang K, Levine AT, Tsetsenis T, Jenson D, Cao F, et al. Dopamine Regulates Aversive Contextual Learning and Associated In Vivo Synaptic Plasticity in the Hippocampus. Cell Rep. 2016;14:1930–9.

[44] Zhang H, Wei W, Zhao M, Ma L, Jiang X, Pei H, et al. Interaction between Abeta and Tau in the Pathogenesis of Alzheimer’s Disease. Int J Biol Sci. 2021;17:2181–92.

[45] Busche MA, Wegmann S, Dujardin S, Commins C, Schiantarelli J, Klickstein N, et al. Tau impairs neural circuits, dominating amyloid-beta effects, in Alzheimer models in vivo. Nat Neurosci. 2019;22:57–64.

[46] Ittner A, Chua SW, Bertz J, Volkerling A, van der Hoven J, Gladbach A, et al. Site-specific phosphorylation of tau inhibits amyloid-β toxicity in Alzheimer’s mice. Science. 2016;354:904–8.

